# The VanS sensor histidine kinase from type-B VRE recognizes vancomycin directly

**DOI:** 10.1101/2023.07.09.548278

**Authors:** Lina J. Maciunas, Photis Rotsides, Elizabeth J. D’Lauro, Samantha Brady, Joris Beld, Patrick J. Loll

## Abstract

<u>V</u>ancomycin-<u>r</u>esistant <u>e</u>nterococci (VRE) are among the most common causes of nosocomial infections and have been prioritized as targets for new therapeutic development. Many genetically distinct types of VRE have been identified; however, they all share a common suite of resistance genes that function together to confer resistance to vancomycin. Expression of the resistance phenotype is controlled by the VanRS two-component system. This system senses the presence of the antibiotic, and responds by initiating transcription of resistance genes. VanS is a transmembrane sensor histidine kinase, and plays a fundamental role in antibiotic resistance by detecting vancomycin or its effects; it then transduces this signal to the VanR transcription factor, thereby alerting the organism to the presence of the antibiotic. Despite the critical role played by VanS, fundamental questions remain about its function, and in particular about how it senses vancomycin. Here, we focus on a purified VanRS system from one of the most clinically prevalent forms of VRE, type B. We show that in a native-like membrane environment, the autokinase activity of type-B VanS is strongly stimulated by vancomycin. We additionally demonstrate that this effect is mediated by a direct physical interaction between the antibiotic and the type-B VanS protein, and localize the interacting region to the protein’s periplasmic domain. This represents the first time that a direct sensing mechanism has been confirmed for any VanS protein.

**Significance Statement:** When <u>v</u>ancomycin-<u>r</u>esistant <u>e</u>nterococci (VRE) sense the presence of vancomycin, they remodel their cell walls to block antibiotic binding. This resistance phenotype is controlled by the VanS protein, a histidine kinase that senses the antibiotic or its effects and signals for transcription of resistance genes. However, the mechanism by which VanS detects the antibiotic has remained unclear, with no consensus emerging as to whether the protein interacts directly with vancomycin, or instead detects some downstream consequence of vancomycin’s action. Here, we show that for one of the most clinically relevant types of VRE, type B, VanS is activated by direct binding of the antibiotic. Such mechanistic insights will likely prove useful in circumventing vancomycin resistance.

## INTRODUCTION

Antibiotic resistance is a global health problem. Pathogenic bacteria are becoming resistant to currently available antibiotics at an alarming rate, while the development of new antibiotics remains slow (1). Important examples of antibiotic-resistant bacteria include the <u>v</u>ancomycin-resistant <u>e</u>nterococci (VRE), which have been identified by the World Health Organization as top priorities for new therapeutic development (2,3). VRE are among the so-called ESKAPE pathogens; these organisms are leading causes of hospital-acquired infections, and few therapeutic options remain for their treatment (4,5).

Ten different VRE genotypes are currently recognized, and are distributed across multiple species of enterococci. These resistance types are denoted as types A, B, C, D, E, G, L, M, N, and P (also known as VanA, VanB, VanC, etc. (6)); currently, the types most commonly associated with human disease are types A and B (7). In all cases, VRE evade vancomycin’s antibiotic action by remodeling their cell wall, preventing the antibiotic from binding its target (8). This mode of resistance requires the acquisition of five key genes, *vanRSHAX*, which are necessary and sufficient to confer vancomycin resistance (9). VanHAX are enzymes that remodel vancomycin’s cell-wall target, and their expression is controlled by a two-component system comprising VanR and VanS. VanR is a transcription factor that is activated by phosphorylation; VanS is a membrane-bound sensor histidine kinase that detects vancomycin and modulates the phosphorylation state of VanR accordingly (10).

No structure is currently known for any full-length VanS protein. However, predictions from sources such as AlphaFold, along with structural information from other histidine kinases, suggest that VanS proteins consist of four structural domains: a sensor domain, a membrane-proximal domain, a <u>d</u>imerization-and-<u>h</u>istidine <u>p</u>hosphotransfer (DHp) domain, and a <u>c</u>atalytic and <u>A</u>TP-binding (CA) domain (11,12) (Figure 1A). The sensor domain is predicted to contain two transmembrane helices flanking a periplasmic region (13,14), and is likely responsible for recognizing the presence of vancomycin. When VanS senses vancomycin, the signal is transduced to the cytoplasmic domains, which coordinate an enzymatic response involving autophosphorylation on a conserved histidine residue, followed by transfer of the phosphoryl group to VanR (9,15). Once phosphorylated, VanR initiates transcription of the resistance genes (16). In the absence of an activating signal, VanS can dephosphorylate VanR, terminating the signaling cascade (17,18). Interrupting this signaling process represents an obvious therapeutic avenue that could salvage the utility of the antibiotic. This goal could be achieved by inhibiting autophosphorylation and phosphotransfer, and/or by stimulating VanR dephosphorylation.

**Figure 1.**
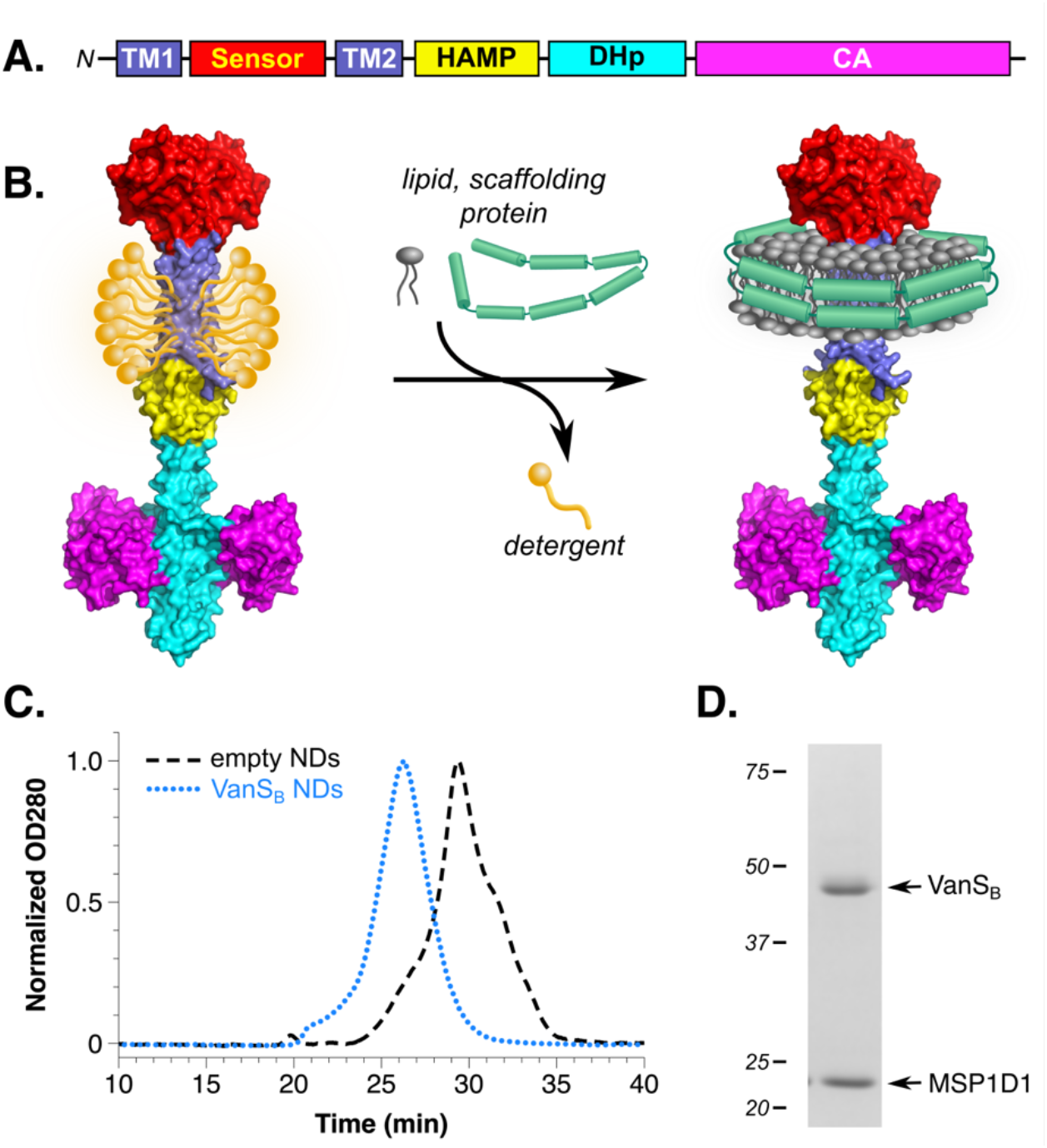
Reconstitution of VanS_B_ into nanodiscs. (*A*) Schematic representation of the typical VanS architecture. Two transmembrane helices (TM1 & TM2) flank a periplasmic sensor domain, the length of which varies significantly among different VanS proteins. The cytoplasmic portion of the protein contains a membrane-proximal region, which typically forms a HAMP domain, followed by the DHp and CA domains. (*B*) For nanodisc formation, detergent-solubilized protein is incubated with lipids and scaffolding protein, after which detergent is removed, triggering the spontaneous formation of protein-belted lipid discs around the hydrophobic face of the protein. The protein model shown represents the structure of full-length VanS_B_, as predicted by AlphaFold; domain colors match those found in panel (*A*) (72). (*C*) Size-exclusion chromatograms reveal that nanodiscs (NDs) containing VanS_B_ elute earlier than empty nanodiscs formed under identical conditions, indicating size increases consistent with the incorporation of the VanS protein. (*D*) Coomassie-stained denaturing SDS-PAGE gels showing a purified VanS_B_ nanodisc preparation, indicating that this preparation contains approximately equimolar amounts of VanS_B_ and the scaffolding protein MSP1D1.

The largest gap in our understanding of vancomycin resistance centers on how VanS recognizes vancomycin. In principle, VanS could detect vancomycin either directly, via a physical interaction with the antibiotic, or indirectly, by sensing a downstream consequence of vancomycin’s action. Multiple different VanS proteins have been studied, including proteins from VRE and homologs from non-enterococcal species, and arguments have been advanced in favor of both models. Chimeric-protein studies, photolabeling, and spectroscopic experiments have all provided data that support a direct-interaction model (19-24); in contrast, an indirect model is supported by the observation that some VanS proteins can be promiscuously activated by many different inhibitors of cell-wall biosynthesis (based on the reasoning that a single binding site on VanS would be unlikely to recognize a wide array of structurally dissimilar effectors) (25-28). Importantly, the direct and indirect models are not necessarily mutually exclusive, nor must a single model apply to all VanS proteins. Thus, considerable uncertainty surrounds the molecular mechanism by which VanS recognizes and transduces the vancomycin signal.

To elucidate this process, we have chosen to examine vancomycin recognition in type-B VRE, focusing on the VanS protein from these organisms (denoted herein as VanS_B_). We purified full-length VanS_B_ and inserted it into nanodiscs. This approach provides the molecular insights gained by reconstituting a signaling system from purified components, while simultaneously maintaining a native-like membrane environment. We investigated the relationship of vancomycin binding to enzymatic activity, showing that vancomycin markedly stimulates VanS_B_ activity, and further demonstrated a direct physical interaction between the VanS_B_ sensor domain and vancomycin. These results link activity changes with direct binding of the antibiotic for the first time for any VanS protein.

## RESULTS

### Reconstitution of VanS into nanodiscs

Membrane proteins are typically purified in detergent-solubilized form, but detergents can reduce stability and/or block access to interaction sites (29-31). The latter effect is of particular concern in the VanRS system, since it has been suggested that vancomycin interacts with VanS at a site directly adjacent to the membrane (22). To provide a more native-like environment for VanS_B_, we reconstituted it into nanodiscs, lipid-bilayer discs belted by an amphipathic scaffolding protein (32,33). VanS_B_ nanodiscs were created by mixing detergent-solubilized protein with *E. coli* lipids and the scaffolding protein MSP1D1, after which detergent was removed, allowing the nanodiscs to spontaneously assemble (Figure 1B). VanS_B_-containing nanodiscs were separated from empty nanodiscs by capturing the His-tagged VanS_B_ protein via nickel-affinity chromatography, after which aggregated material was removed on a size-exclusion column. Re-injection of the purified material gave symmetrical peaks eluting significantly earlier than empty nanodiscs (Figure 1C). SDS-PAGE analysis revealed that VanS_B_ and MSP1D1 are present in the nanodiscs in an approximate 1:1 stoichiometry (Figure 1D). Since each disc is known to contain two copies of the MSP1D1 scaffolding protein (33), this implies that on average, each nanodisc contains two copies of VanS_B_. Since sensor histidine kinases typically form obligate homodimers (34), each nanodisc likely contains a single homodimer.

Only a small number of sensor histidine kinases have been reconstituted into nanodiscs (35-42), and this is the first report of this approach being applied to any VanS protein. These reconstituted VanS_B_ protein preparations provide the foundation for the *in-vitro* experiments described below, and allow us to avoid the complications associated with the use of detergent.

### VanS_B_ is active after reconstitution into nanodiscs

Sensor histidine kinases autophosphorylate in response to signals; they also modulate the phosphorylation states of their response regulators, transferring their phosphoryl group to the response regulator. Some sensor histidine kinases exert additional control via a phosphatase activity that dephosphorylates their response regulators in the absence of a signal (34). These three activities reside in the cytosolic domains of the kinases, and are regulated by the transmembrane and sensor regions in response to stimulus detection. In the case of VanS_B_, a construct containing only the cytosolic domain has been shown to display all three enzymatic activities (17). To examine the behavior of full-length VanS_B_ in a nanodisc environment, we first focused on the autophosphorylation function. We monitored activity using a modified Western blot assay (Figure 2A), in which VanS_B_ was allowed to autophosphorylate with ATPγS, after which the thiophosphohistidine was alkylated and detected by antibody (43). Like VanS_A_, the VanS protein found in type-A VRE (44), VanS_B_ is strongly sensitive to the specific membrane mimetic used, and shows essentially no activity in the presence of the detergents dodecyl maltoside (DDM) or lauryl dimethyl amine oxide (LDAO), and modest activity in dodecyloctapolyoxyethylene (C_12_E_8_). The nanodisc preparation of VanS_B_ is autokinase-active, validating the use of this approach. Interestingly, however, reconstitution of VanS_B_ into nanodiscs leads to only modest basal levels of autokinase activity, failing to surpass the levels seen in the presence of C_12_E_8_.

**Figure 2.**
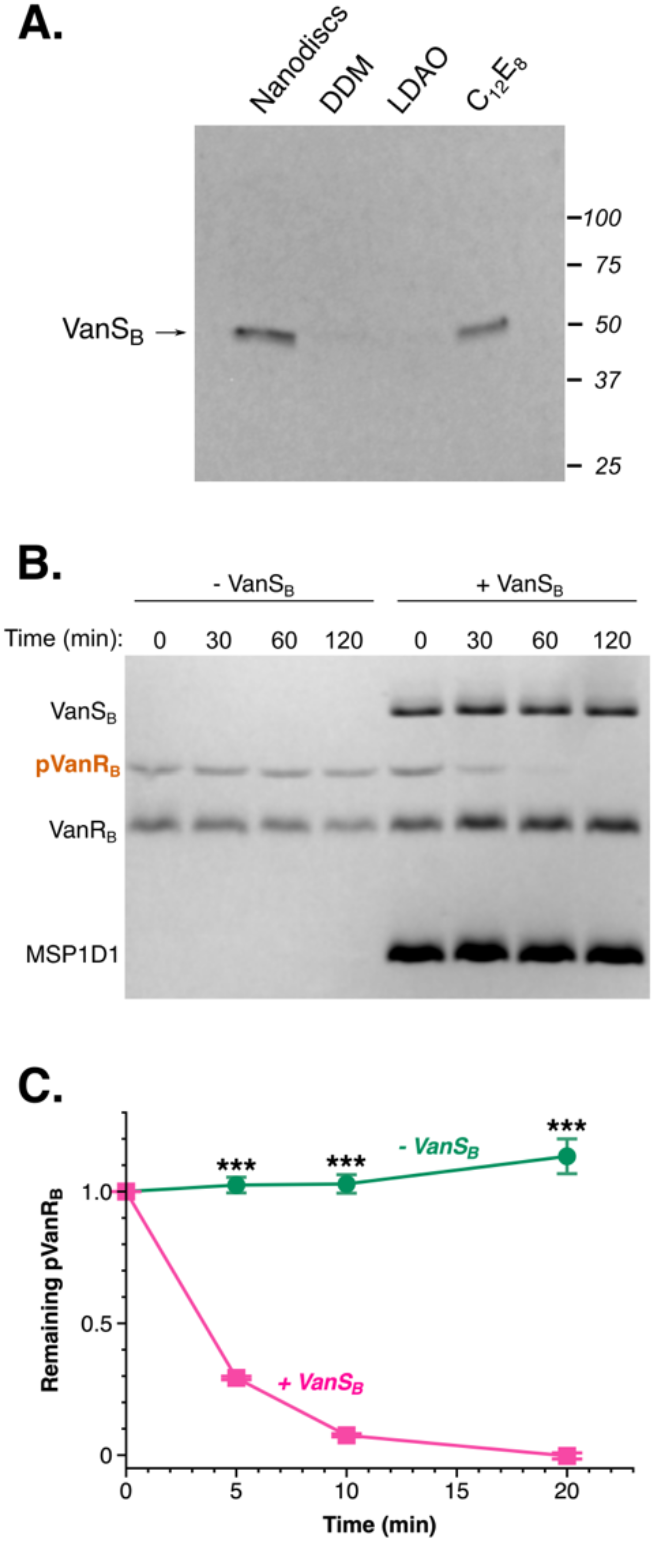
Enzymatic activities of purified VanS_B_. (*A*) VanS_B_ activity is strongly sensitive to the membrane-mimetic environment. In each reaction, 0.3 µM VanS_B_ was incubated with 1 mM ATPγS for 30 minutes. The thiophosphohistidine was then alkylated with *p*-nitrobenzyl mesylate (PNBM), which was subsequently detected by Western blot. Equal amounts of protein are loaded in each lane. VanS_B_ shows very little activity in the presence of the detergents DDM and LDAO, and modest activity in C_12_E_8_ and nanodiscs. (*B*) & (C) Dephosphorylation activity of VanS_B_. Phosphorylated VanR_B_ was incubated ± VanS_B_-containing nanodiscs, and the time-dependent decrease in phosphorylation was monitored using Phos-Tag™ gels. Panel (*B*) shows representative gel, and panel (*C*) shows the quantitation (*n* = 3). Asterisks represent *p*-values calculated using an unpaired T-test; ***, *p*-value < 0.0001. This experiment illustrates the high intrinsic stability of phospho-VanR_B_ (17), consistent with the observation that VanR_B_ can be isolated in a partially phosphorylated form after expression in *E. coli* (SI Appendix, Fig. S8).

We next probed dephosphorylation activity, incubating VanS_B_ with the phosphorylated form of its cognate VanR protein, and quantifying the time-dependent decrease in phospho-VanR using Phos-tag™ gels (45,46). As anticipated, VanS_B_ was found to catalyze the dephosphorylation of VanR_B_ (Figure 2B, 2C).

We did not quantify phosphotransfer activity, since the catalytic machinery mediating this process typically resides primarily in the response regulator, rather than the sensor kinase (47). We did confirm, however, that VanS_B_ is capable of phosphotransfer to its cognate VanR_B_ partner (SI Appendix, Fig. S1). Hence, our reconstituted nanodisc preparation of VanS_B_ possesses all three relevant enzymatic activities.

### Vancomycin’s effect on VanS_B_ activity

We hypothesized that if vancomycin directly activates VanS_B_, then the antibiotic should increase the enzyme’s autokinase activity and/or decrease its phosphatase activity; either effect would tend to increase cellular levels of phospho-VanR_B_. We first investigated the effect of vancomycin on the autokinase activity of VanS_B_, and found that it stimulated autophosphorylation in a concentration-dependent manner (Figure 3A-C). We then showed that this stimulatory effect was also observed using ^32^P-ATP as a kinase substrate, confirming that this result was not unique to ATPγS (SI Appendix, Fig. S2). In addition, we prepared a VanS_B_ construct lacking the C-terminal His-tag and demonstrated that its autophosphorylation activity is also stimulated by vancomycin, to a similar extent as that seen in the tagged protein, confirming that the affinity tag is not interfering with the protein’s function (SI Appendix, Fig. S3).

**Figure 3.**
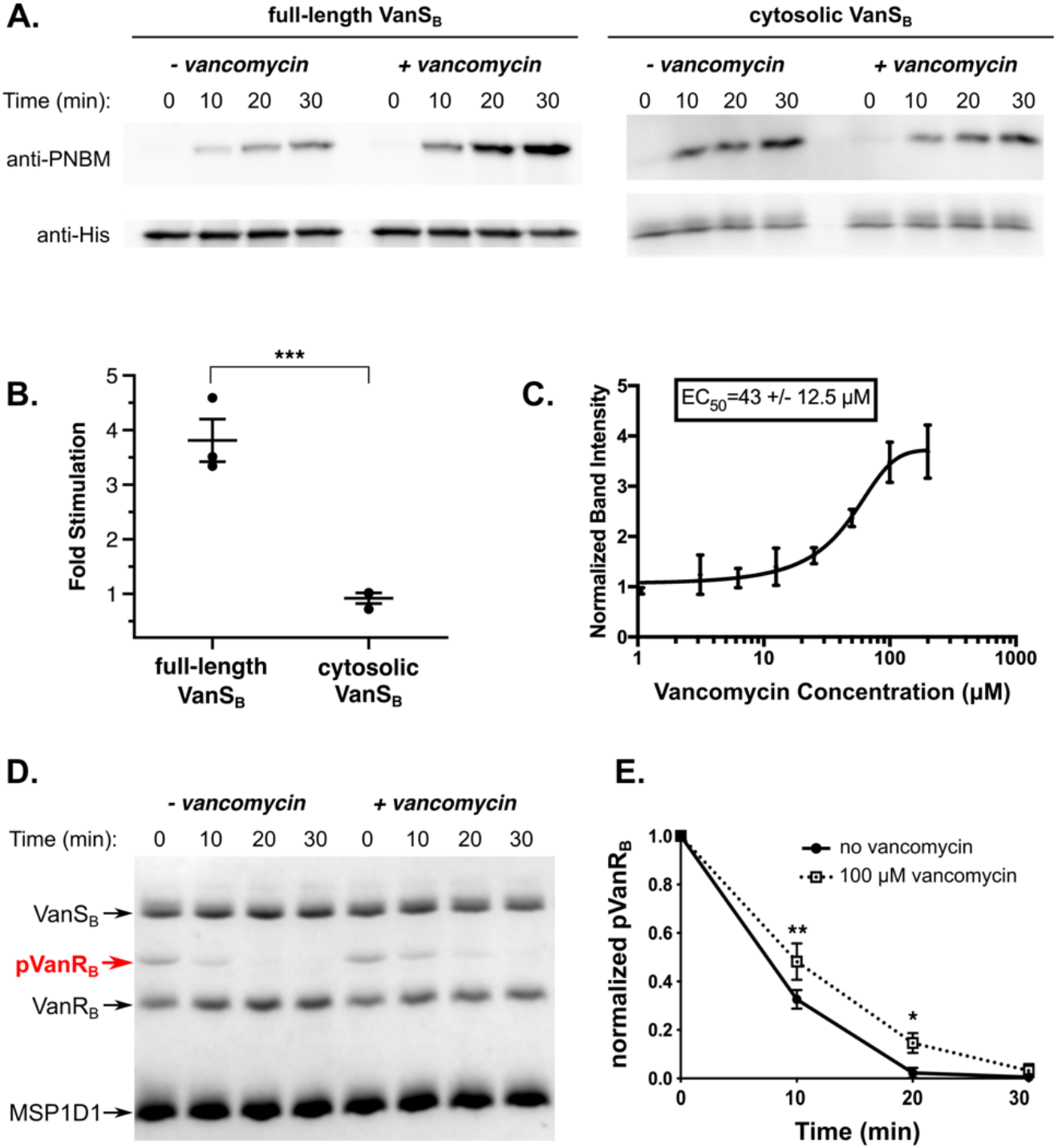
Effect of vancomycin on VanS_B_ activities. *(A*) Representative Western blots showing an autophosphorylation time course for full-length VanS_B_ (reconstituted in nanodiscs) and its cytosolic domain, ± 100 µM vancomycin. (*B*) Quantitation of vancomycin stimulation of autophosphorylation activity (*n* = 3). Shown is the fold increase in autophosphorylation levels (relative to no antibiotic) after 30-min treatment with 100 µM vancomycin. ***, *p*-value < 0.005 (*C*) Dose dependence for vancomycin stimulation of VanS_B_ autophosphorylation activity. (*D*) A representative Phos-tag™ gel demonstrating that 100 µM vancomycin has a modest inhibitory effect on the dephosphorylation activity of VanS_B_. (*G*) Quantitation of three dephosphorylation experiments for VanS_B_. *, *p*-value < 0.05; **, *p*-value < 0.01.

When VanS_B_ is embedded in a nanodisc, both its extracellular and intracellular domains will be accessible to a water-soluble molecule such as vancomycin; however, in the native cellular environment, only the periplasmic domain will be accessible to the antibiotic. Therefore, any effect that vancomycin has on the enzyme’s activity should be mediated through the periplasmic sensor domain, and not through any of the cytosolic domains. To confirm the absence of any such nonspecific effects, we purified a cytosolic VanS_B_ construct lacking the sensor domain, and showed that vancomycin has no effect on its autophosphorylation activity (Figure 3B). Thus, the stimulatory effect of vancomycin is specific to the full-length VanS_B_ protein.

Finally, we tested whether vancomycin influences the dephosphorylation activity of VanS_B_, and found that it slightly decreased the enzyme’s dephosphorylation of VanR_B_ (Figure 3D-E). In summary, vancomycin increases VanS_B_’s autophosphorylation activity and modestly decreases its phosphatase activity; both effects are consistent with vancomycin directly stimulating VanRS signaling in type-B VRE.

### Vancomycin binds to the periplasmic sensor domain of VanS_B_

Vancomycin’s effect on the enzymatic activity of purified VanS_B_ suggested that the antibiotic might bind directly to the enzyme. We reasoned that the likely site of binding is the protein’s periplasmic region, as it is the only portion of the protein accessible from the outside of the cell. The VanS_B_ periplasmic domain contains roughly 100 residues, and it is predicted to adopt a PAS-domain fold (SI Appendix, Fig. S4) (48,49). To test whether the isolated periplasmic domain is capable of binding the antibiotic, we prepared expression constructs for the appropriate region, corresponding to VanS_B_ residues 31-132. We produced two constructs, the first encoding a single copy of the periplasmic sensor, and the second containing two copies of the sensor in tandem, separated by a short linker (Figure 4A). The latter construct was designed to encourage dimerization, since full-length sensor histidine kinases are thought to be obligate dimers (34). Both proteins were purified to homogeneity (SI Appendix, Fig. S4A).

**Figure 4.**
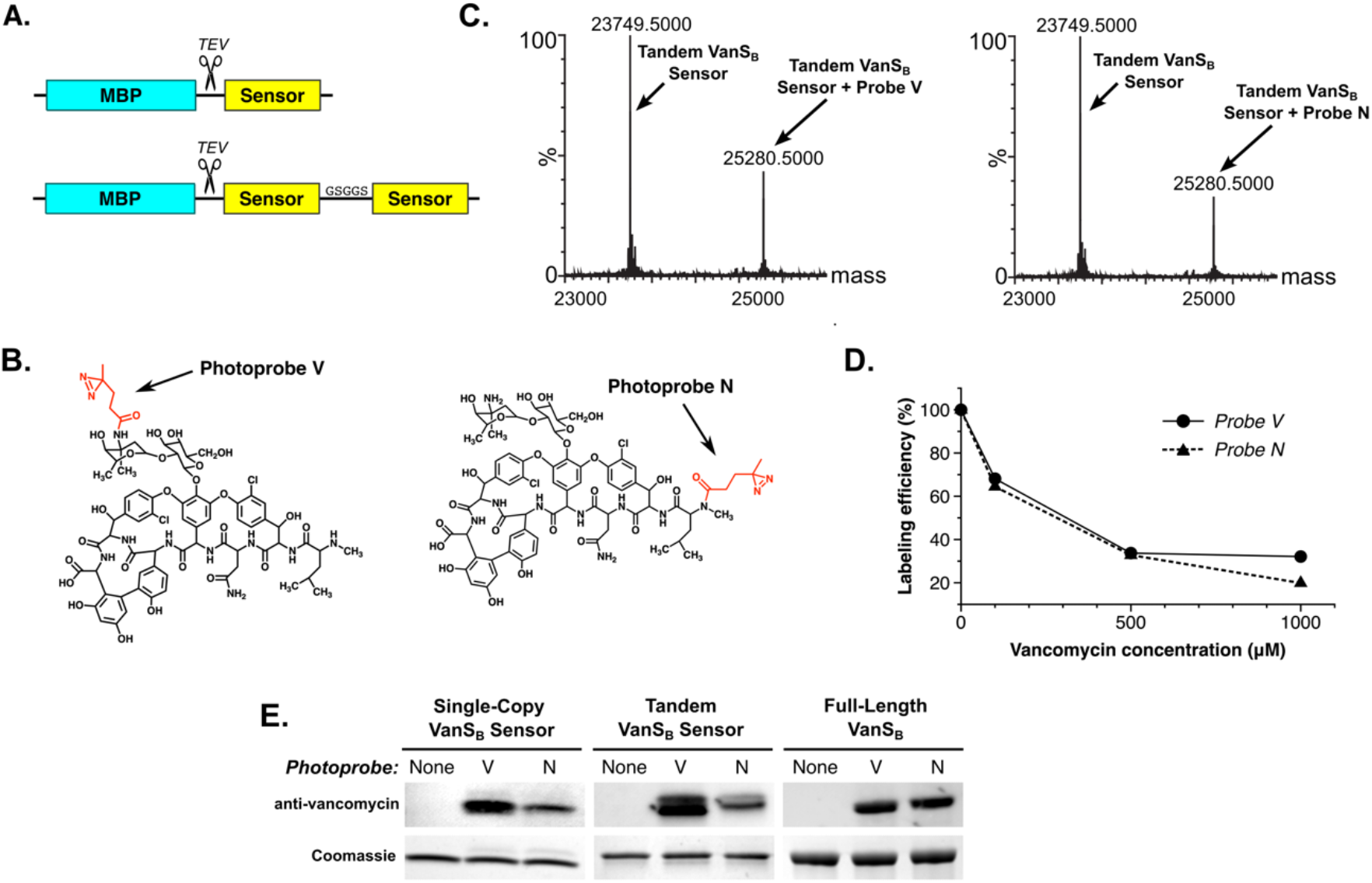
The VanS_B_ periplasmic sensor domain can be labeled with vancomycin-based photoprobes. (*A*) The VanS_B_ periplasmic sensor domain (residues 31-132) was expressed as a cleavable fusion with MBP, in both single-copy and tandem forms, and purified to homogeneity (SI Appendix, Fig. S4). (B) Two photoprobes were used, in which a photo-active diazirine was attached to either the antibiotic’s vancosamine sugar (Photoprobe V) or its N-terminus (Photoprobe N). (C) Mass spectrometric evidence that both photoprobes label the purified tandem sensor-domain construct. (D) Vancomycin competes with both photoprobes for binding to the tandem sensor domain. (E) Confirmation of photolabeling using an anti-vancomycin Western-blot assay, showing labeling of both the isolated single-copy and tandem sensor domains, as well as full-length VanS_B_. Coomassie-stained gels serve as loading controls. Photoprobe N typically labels VanS_B_ constructs less efficiently than Photoprobe V, as judged by both Western blotting and mass spectrometry (see panel (C)); this may reflect differences in the diazirine group’s proximity to the protein, as determined by the conformation of the bound antibiotic.

We first probed antibiotic binding to the periplasmic-domain constructs using two novel photoaffinity probes, in which a diazirine photolabel is attached at either end of the vancomycin molecule (50). Specifically, the photolabel is attached to either the antibiotic’s vancosamine sugar (Probe V) or its N-terminus (Probe N; see Figure 4B). Both photoprobes efficiently labeled the single-copy and tandem VanS_B_ sensor proteins, as demonstrated by mass spectrometry and anti-vancomycin Western blotting (Figure 4C & 4E). No evidence was seen for double labeling of the tandem sensor domain, even though in principle it can contain two distinct ligand-binding sites (51); however, a covalent dimer is not required for the interaction, since photolabeling was also observed for the single-copy construct. Unlabeled vancomycin competed effectively with both photoprobes, indicating that the protein-photoprobe interaction is specific (Figure 4D). Notably, both photoprobes also label the full-length VanS_B_ protein, indicating that the observed interaction with the sensor domain is not an artifact caused by removing the periplasmic domain from the context of the intact protein molecule. Overall, these results provided strong evidence of a specific, direct interaction between vancomycin and the VanS_B_ protein.

We next sought to quantify this binding interaction, using fluorescence anisotropy to assess recognition of BODIPY-FL-labeled vancomycin (Figure 5). The single-copy VanS_B_ sensor domain was found to bind to the labeled vancomycin with a dissociation constant (*K*_D_) of 20.3 μM, while the tandem sensor bound with a *K*_D_ of 10.7 µM; the fact that the single-domain *K*_D_ value is almost exactly twice that of the tandem domain suggests that there is no cooperativity between the two domains, at least in the constructs used. We then generated negative controls, creating single- and tandem-domain versions of the periplasmic sensor domain from the *B. subtilis* histidine kinase PhoR. The PhoR sensor domain adopts a PAS fold, similar to that predicted for the VanS_B_ periplasmic domain (52). However, PhoR is not expected to bind to vancomycin, and indeed, neither PhoR construct bound to BODIPY-FL-vancomycin (Figure 5B). Next, to test whether the binding observed in the anisotropy experiment was fluorophore-dependent, we created a new fluorescent probe by labeling vancomycin with AlexaFluor488, instead of BODIPY-FL (SI Appendix, Fig. S5). The fluorophores are attached at the same position in both probes, but differ significantly in structure and hydrophobicity. The tandem VanS_B_ sensor protein bound to AF488-vancomycin with a *K*_D_ value of 11.0 μM, essentially identical to the affinity measured for BODIPY-FL-vancomycin, allowing us to rule out any nonspecific interactions between the sensor domain and the fluorophore.

**Figure 5.**
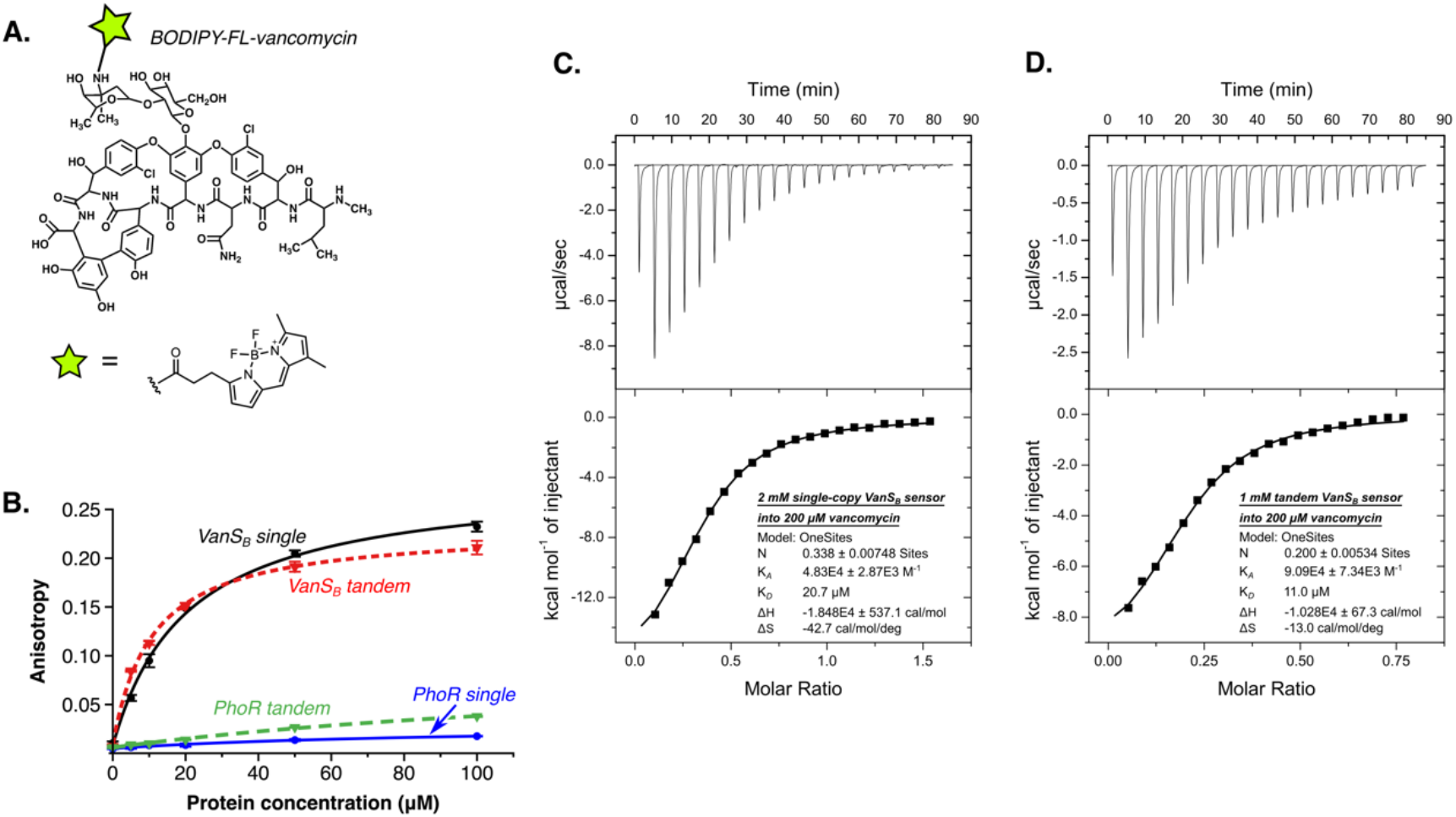
Vancomycin binds directly to the periplasmic sensor domain of VanS_B_. (*A*) Fluorescently labeled vancomycin derivative used for fluorescence anisotropy measurements. (*B*) BODIPY-FL-vancomycin binds to both the single and tandem sensor-domain constructs of VanS_B_. In contrast, single and tandem constructs of the PhoR negative control fail to bind the antibiotic. (*C*) & (*D*), Isothermal titration calorimetry confirms vancomycin binding by the single-copy (*C*) and tandem (*D*) VanS_B_ sensor-domain constructs. The related glycopeptide antibiotic teicoplanin fails to bind to the sensor domain (SI Appendix, Fig. S5), which is consistent with the known behavior of type-B VRE.

Finally, we used isothermal titration <u>c</u>alorimetry (ITC) to measure the binding of label-free vancomycin to the VanS_B_ sensor domain, obtaining *K*_D_ values of 20.7 μM for the single-copy sensor domain and 11.0 μM for the tandem sensor, in excellent agreement with the results of the fluorescence-anisotropy experiments (Figure 5C & 5D). Importantly, the related glycopeptide antibiotic teicoplanin does not bind the VanS_B_ sensor (SI Appendix, Fig. S6), consistent with observation that teicoplanin does not induce resistance in type-B VRE (26). Taken together, these results clearly demonstrate a direct and specific binding interaction between vancomycin and the periplasmic sensor domain of VanS_B_.

## Discussion

Of the various organisms that comprise VRE, those displaying type-B resistance are perhaps the most rapidly growing threat. While at least ten different VRE genotypes are known, infections in humans are predominantly caused by either types A or B, perhaps because both of these resistance phenotypes are associated with mobile genetic elements (53). Historically, type A has proven more prevalent than type B, but the incidence of the latter has been steadily rising (54-58). Type-B resistance confers high-level resistance against vancomycin, but not against related glycopeptide antibiotics (59). It is principally found in *E. faecium*, but has also been observed in other enterococcal species; in addition, non-enterococcal species, including gut anaerobes such as *Clostridioides* spp., can contain *vanB* genes, serving as reservoirs from which these genes can ultimately be transferred to enterococci (60).

The mechanisms by which any VRE evade vancomycin remain incompletely understood, with the largest open questions being how VanS recognizes the antibiotic and transduces this signal. Answers to these questions will likely inform the development of new therapeutic avenues. To date, most effort has been devoted to studying VanS proteins from type-A VRE and from non-enterococcal organisms, and these studies have not yet conclusively determined whether these proteins are activated by direct or indirect mechanisms. In contrast, the full-length VanS_B_ protein had not been well characterized prior to this work. Here, we present definitive evidence for regulation of VanS_B_ activity via a direct interaction with vancomycin. Specifically, vancomycin increases VanS_B_’s autophosphorylation activity and slightly decreases its phosphatase activity; both effects will lead to increased levels of phospho-VanR_B_, thereby stimulating the transcription of the resistance operon. We further provide evidence of a direct physical interaction between vancomycin and VanS_B_. This is the first time that direct binding of the antibiotic has been linked to activity changes in a VanS protein.

While we observe clear evidence for vancomycin binding to VanS_B_, the binding affinities and EC_50_ value for autokinase stimulation are significantly higher than the ∼2 µM vancomycin concentration reported to induce transcription of resistance genes in type-B VRE (26). There are many potential explanations for this discrepancy. Different VanB isolates vary widely in their susceptibility to vancomycin (61), and it is possible that our chosen VanS_B_ sequence simply represents a relatively low-sensitivity vancomycin binder. It may also be true that our simplified *in-vitro* assay system, while useful, is not a perfect mimic of the environment found *in vivo*. For example, our assay system contains no peptidoglycan or lipid II; in bacterial cells, both of these components would tend to recruit vancomycin to the vicinity of VanS_B_’s membrane-proximal sensor domain, giving rise to a high local concentration of the antibiotic. In particular, lipid II levels increase in response to vancomycin activity (62), which could directly modulate VanS_B_ activity. Another factor to consider is that our nanodiscs contain, on average, only two copies of the VanS_B_ molecule per disc, likely forming a homodimer; if a higher-order assembly of dimers is required for complete activation (35), this could not be achieved in our system. Alternatively, some other protein (encoded in either the *vanB* operon or the enterococcal genome) might be required to achieve maximum sensitivity. Finally, we note that our nanodisc preparations use commercially available *E. coli* lipids, in which phosphatidylethanolamine is the predominant lipid species, along with smaller amounts of phosphatidylglycerol and cardiolipin (63). In contrast, membranes of enterococci feature primarily phosphatidylglycerol and cardiolipin (64), and this difference in lipid environment could conceivably affect VanS_B_ function.

Different VanS proteins perform equivalent functions in their respective VRE strains, but may do so by different mechanisms. The key to any mechanistic differences is likely to reside in the various proteins’ sensor domains. For example, while VanS_A_ and VanS_B_ share ∼19% sequence identity and 36% overall similarity, this similarity is most pronounced in the cytoplasmic regions of these proteins, whereas the periplasmic sensor domains are highly dissimilar in both size and sequence (SI Appendix, Fig. S7). VanS_A_ is typical of most VanS proteins in that it contains a short periplasmic domain—in this case, ∼37 residues. In contrast, the VanS_B_ periplasmic domain is ∼101 residues in length. The VanS_B_ periplasmic region is predicted to adopt a PAS domain fold; this appears to be unique to type-B VanS proteins, as the periplasmic regions of VanS proteins from other VRE types are too short to accommodate such a structure (SI Appendix, Fig. S8). In fact, phylogenetic analysis reveals that VanS genes from type-B VRE form an outlier cluster, distinct from the VanS genes associated with other VRE types (65). Hence, the sensor domain that allows VanS_B_ to bind directly to vancomycin is evolutionarily distinct, making it difficult to extrapolate from the type-B example and draw mechanistic inferences about VanS proteins from other VRE types.

The elucidation of the VanS_B_ sensing mechanism has important translational implications. For example, the confirmation of a direct binding interaction between the antibiotic and the protein rationalizes why type-B VRE are only resistant to vancomycin, and not to other glycopeptide antibiotics that share vancomycin’s mechanism of action, but differ in structure. This holds out the possibility of generating new “stealth antibiotics,” vancomycin derivatives that are not recognized by VanS_B_. Alternately, one can envision small molecules that block vancomycin binding to VanS_B_, acting as antibiotic adjuvants that restore vancomycin efficacy by suppressing VanRS signaling.

## Materials and Methods

### Expression constructs

Primers used for subcloning are shown in Table S1. pMSP1D1 was obtained from Addgene (catalog #20061). The genes for full-length VanS_B_ and VanR_B_ (Uniprot IDs Q47745 and Q47744, respectively) were codon-optimized for expression in *E. coli* cells by Genscript (Piscataway, NJ) and subcloned into the in-house pETCH vector, producing expression constructs containing C-terminal His_6_ tags (66). To produce a tagless version of VanS_B_, the gene was inserted into the in-house pETHSUL vector, thereby fusing a cleavable His_6_-SUMO sequence to the N-terminus of VanS_B_ (66). The cytosolic form of VanS_B_ (cVanS_B_) was previously determined to comprise residues M159-L447 (17), and the corresponding gene fragment was subcloned into pETCH. The periplasmic domain of VanS_B_ is located between the two transmembrane helices, which are predicted to consist of residues 10-28 and 135-155 (67). Therefore, to produce a tandem periplasmic-domain construct, two gene fragments were designed. Both encoded residues Q31 to G132 of VanSB, and also included an overlapping linker region encoding the linker sequence GSGGS connecting the C-terminus of the first copy to the N-terminus of the second copy. These fragments were assembled using the NEBuilder HiFi Assembly kit into pMal-c6T (NEB #N0378S). The resulting construct encodes a TEV-cleavable 6xHis-MBP tag fused to two copies of the VanS_B_ periplasmic domain. The single-copy construct was created by introducing the sequence TAAT via site-directed mutagenesis, placing an enhanced stop codon after the first copy of the VanS_B_ sensor domain (68). A similar strategy was used to produce tandem and single-copy constructs of the *B. subtilis* PhoR periplasmic domain (Uniprot P23545 aa 32-150), amplifying the appropriate region from *Bacillus subtilis* subsp. *subtilis* strain 168 (ATCC cat. no. 23857).

### Protein expression and purification

C-terminally His-tagged VanS_B_ was purified following a protocol previously described for VanS_A_ (44), with minor modifications. Briefly, pelleted cells from 6-L cultures were resuspended in IMAC-A buffer (40 mM Tris pH 8, 300 mM NaCl, 25 mM imidazole) containing EDTA-free protease inhibitor tablets (Thermo-Pierce) and then lysed in a C5 Emulsiflex cell homogenizer at 10,000-15,000 psi. Membranes were isolated by centrifugation and homogenized in IMAC-A buffer, after which proteins were solubilized by the addition of 1% (w/v) *N*-dodecyl-β-D-maltopyranoside (DDM) for 1 hour at 4°C. The solubilized membranes were centrifuged at 150K *g* for 50 min, and the resulting supernatant was syringe-filtered (0.45 μm) and loaded onto a 1-mL IMAC-HP column (Cytiva) equilibrated in IMAC-A buffer containing 0.2% DDM. Protein was eluted with a gradient from 25 to 350 mM imidazole.

To produce the tagless version of VanS_B,_ the pETHSUL-VanS_B_ plasmid was transformed into BL21(DE3) pLysS cells. Cells were grown in TB media with 100 μg/mL ampicillin and 34 μg/mL chloramphenicol to an OD600 of 0.8, after which the temperature was reduced to 16 °C. Once the OD600 reached 1.2, 1 mM IPTG was added, and the cultures were shaken for 22 hours, after which cells were harvested by centrifugation and frozen at -80 °C. After thawing, cells were lysed and the His_6_-SUMO-VanS_B_ protein was purified following the identical procedure used for the C-terminally His-tagged VanS_B_ protein.

VanR_B_ was co-expressed with GroEL/S chaperones. pETCH-VanR_B_ and pGro7 (Takara Cat# 3340) were cotransformed into BL21(DE3) cells. Overnight cultures were diluted 1:500 in LB media containing 100 μg/mL ampicillin and 34 μg/mL chloramphenicol. Immediately after inoculation, arabinose (1 mg/mL) was added to induce chaperone expression. At OD600 = 0.5, the temperature was lowered to 30°C, IPTG was added (0.2 mM), and cells were incubated for 4 hours. Cells harvested from 3-L cultures were resuspended in 40 mM Tris pH 8, 500 mM NaCl, 25 mM imidazole, 10% w/v glycerol, 5 mM MgCl_2_ (Buffer IMAC-A2) containing 10 μg/mL DNase, 2 μg/mL RNase, and EDTA-free protease inhibitor tablets. The resuspended cells were lysed by homogenization, after which Triton-X100 was added to the lysate at a final concentration of 0.1% (v/v) and the sample incubated at 4 °C for 1 hour. The lysate was centrifuged at 150K *g* for 1 hour, filtered (0.45-μm), and loaded onto a 1-mL IMAC-HP column equilibrated in IMAC-A2 containing 0.1% Triton-X100. The column was washed with 20 CV of IMAC-A2 containing 0.1% Triton-X100, 20 CV of IMAC-A2 without detergent, and 20 CV of 10% IMAC-B2 (40 mM Tris pH 8, 500 mM NaCl, 350 mM imidazole, 10% glycerol, 5 mM MgCl_2_). VanR_B_ was eluted with a gradient from 10-100% IMAC-B2. Fractions containing VanR_B_ were pooled, concentrated to 5 mL, syringe-filtered (0.22-μm), and injected onto a Sephacryl S300 16/60 size-exclusion column (Cytiva) equilibrated in IMAC-A2 buffer lacking imidazole. Aliquots of VanR_B_ were flash-frozen in liquid nitrogen and stored at -80°C.

All periplasmic-domain constructs were expressed in BL21(DE3) cells. Cells were grown in LB broth with 0.2% glucose at 37°C to an OD_600_ of ∼0.6, at which point temperature was reduced to 16°C and IPTG was added to a final concentration of 0.2 mM. Cells were harvested after ∼22 hours of shaking at 225 RPM. Cells from two-liter growths were lysed in buffer containing 20 mM Tris pH 7.5, 200 mM NaCl, 5 mM MgCl_2_, 1mM EDTA, and protease inhibitors (ThermoFisher #A32963). Cell lysate was clarified by centrifugation at 10,000x*g* for 20 minutes then at 117,000x*g* for one hour, after which the soluble supernatant was filtered (0.45 μm) and loaded onto a 42-mL column packed with amylose resin (NEB #E8021L) equilibrated with Buffer A3 (20mM Tris pH 7.5, 100mM NaCl). The column was washed with Buffer A3, then the captured protein was eluted in batch using Buffer A3 supplemented with 50 mM maltose. The eluted protein was mixed with 2 mg of TEV protease and dialyzed overnight against one liter of IEX buffer (20 mM Tris pH 7.5) containing 1 mM DTT. The following day, the dialysate was loaded onto a 5-mL HiTrap Q HP column (Cytiva) equilibrated with IEX buffer. The column was first washed with IEX buffer, followed by IEX buffer supplemented with 100 mM NaCl. At this point, a 5-mL HiTrap IMAC HP column was attached downstream of the Q column and proteins were eluted using IEX buffer containing 600 mM NaCl. The IMAC column served to capture the TEV protease, the cleaved 6xHis-MBP tag, and any uncleaved protein during elution. The eluted protein was concentrated to 10 mL and loaded onto a Sephacryl-S100 26/60 column equilibrated with Buffer A3. Fractions containing the desired proteins were analyzed by SDS-PAGE for purity, pooled, and concentrated to at least 5 mg/mL.

The cytosolic domain of VanS_B_ (cVanS_B_) was purified similarly to VanR_B_ with one modification: 21 mM *N*-decyl-*N,N*-dimethylamine-*N*-oxide (DDAO) was added to the lysate and buffer IMAC-A2. MSP1D1 was expressed and purified as previously described (32,33).

### Nanodisc Assembly and Purification

4 mL of a 25 mg/mL chloroform solution of *E. coli* total lipid extract (Avanti Polar Lipids, Inc.) were placed in a clean glass vial. The chloroform was evaporated with a stream of N_2_ gas and the vial placed in a lyophilizer overnight. The dried lipids were resuspended in 3.75 mL of 4% (w/v) DDM in ultrapure water. The suspension was sonicated for 10 min, freeze-thawed, sonicated for an additional 10 min, and then stored on ice. Lipid concentration was determined by phosphorous assay (69,70). Nanodiscs were assembled by combining VanS, MSP1D1, and *E. coli* lipids in a mole ratio of 1:10:600 in a final volume of 500 μL, with a final VanS concentration of 4 μM. After a 40-min incubation at room temperature, the 500-μL nanodisc mixture was added to 200 μL wet Bio-Beads (Bio-Rad Cat# 1523920) in a fresh 1.5-mL tube. Bio-Beads were prepared as described in (71). The BioBead-containing mixtures were rotated end-over-end at 4°C overnight. The following day, Bio-Beads were removed by filtration and the concentration of imidazole was reduced to 30 mM by diluting the sample with cold ND buffer (20 mM Tris pH 7.4,150 mM NaCl). The sample was syringe-filtered (0.22 μm) and applied to a gravity-flow column containing a 600-μL bed volume of charged nickel resin equilibrated in IMAC A buffer. The column was washed with 4 mL of cold IMAC A and the VanS-containing nanodiscs were eluted with 1.8 mL of cold 1:1 mixture of IMAC A and IMAC B buffers. The eluate was concentrated to 250 μL using a 100-kDa MWCO 0.5-mL concentrator, filtered (0.22 μm), and injected onto a Superdex 200 10/300 column (Cytiva) equilibrated in ND buffer at room temperature. Peak fractions were pooled, concentrated, quantified, and used for activity assays.

To generate nanodiscs containing untagged VanS_B_, assemblies were performed as previously described with the following modifications. A mole ratio of 1:10:700 was used in a final volume of 500 uL, with a His_6_-SUMO-VanS_B_ concentration of 4 uM. The 500-uL nanodisc mixture was added to 300 uL of wet BioBeads. After syringe-filtration (0.22 μm), the sample was applied to a 1-mL IMAC HiTrap HP column equilibrated in IMAC A Buffer. The column was washed with cold IMAC A and the His_6_-SUMO-VanS_B_-containing nanodiscs were eluted with cold IMAC B buffer. The eluate was dialyzed against 2 L of 20 mM Tris pH 7.4, 150 mM NaCl overnight at 4°in a 3K MWCO dialysis cassette (Thermo Slide-A-Lyzer). Prior to dialysis, 1 mg of purified SUMO protease (dtUD1; 66) was added to cleave the His_6_-SUMO tag. The dialysate was then applied to the same IMAC column and the flowthrough was collected. The sample was concentrated to 250 μL using a 100-kDa MWCO 0.5-mL concentrator, filtered (0.22 μm), and injected onto a Superdex 200 10/300 column as described above.

### Autophosphorylation Assays

The autophosphorylation assays using ATP*γ*S were performed as described (44). Briefly, VanS was incubated in 50 mM KCl, 10 mM MgCl_2_, 50 mM Tris pH=7.4 with 1 mM ATP*γ*S for specified times, after which the reaction was quenched by addition of EDTA and PNBM was added to alkylate the thiophosphohistidine. Samples were analyzed by SDS-PAGE and Western blotting, probing with an anti-PNBM antibody (Abcam ab92570). Detailed methods are provided in the Supporting Information.

Autophosphorylation experiments using *γ*−^32^P-labeled ATP (PerkinElmer BLU002Z250UC, 10 mCi/mL) were set up similarly to those using ATP*γ*S. A 50-μL autophosphorylation reaction contained 1X reaction buffer, 2 μL of the ^32^P-ATP stock, 1 mM cold ATP, and VanS_B_-containing nanodiscs at 0.5 μM. Reactions were quenched by the simultaneous addition of EDTA and SDS-PAGE loading buffer and the samples analyzed by SDS-PAGE.

### Dephosphorylation Assays

The VanR_B_ protein as purified from *E. coli* was observed to be partially phosphorylated, presumably as a result of nonspecific phosphorylation by either endogenous histidine kinases or non-enzymatic phosphoryl donors (SI Appendix, Fig. S9). This VanR_B_ preparation was therefore used to assess the dephosphorylation activity of VanS_B_. Partially phosphorylated VanR_B_ protein was incubated at room temperature with VanS_B_; the reaction was quenched at various time points by the addition of SDS-PAGE loading buffer, after which samples were analyzed by electrophoresis on Phos-tag™ gels (Fujifilm Cat. # 195-17991). Additional details are provided in the Supporting Information.

### Photolabeling Reactions

Photoprobes containing diazirine groups attached to either the amino group of vancomycin’s vancosamine sugar (Probe V) or to the antibiotic’s N-terminus (Probe N) were prepared as described (50). Photolabeling was initiated by irradiation with 365-nm light, and samples were analyzed by mass spectrometry and Western blotting; details are provided in the Supporting Information.

### Vancomycin Binding Assays

Fluorescence anisotropy experiments were conducted in 20-μL volumes containing 100 nM BODIPY-FL-vancomycin (ThermoFisher #V34850) or Alexa Fluor 488-vancomycin in 20 mM Tris pH 7.5 buffer containing 100 mM NaCl and 0.01% Triton X-100, using black 384-well small-volume plates (Greiner Bio-One # 784077). Anisotropy measurements were conducted at room temperature using a Tecan Spark microplate reader. Samples were excited at 490 nm and emission read at 525 nm, using a 10 nm bandpass for both emission and excitation. A 510 nm dichroic mirror was used to condition the emitted signal. Single-copy and tandem-copy versions of the VanS_B_ periplasmic domain were used with the BODIPY-FL probe; the tandem-copy protein was used with the AlexaFluor 488 probe to confirm that there was no dependence of binding on the specific identity of the fluorophore.

ITC experiments were carried out using a MicroCal VP-ITC calorimeter (Malvern Panalytical). Protein and vancomycin stock solutions were exhaustively dialyzed against 20 mM Bicine pH 7.5. The sample cell was filled with 200 μM vancomycin and the injection syringe was loaded with 2 mM VanS_B_ single-copy periplasmic-domain protein or 1 mM VanS_B_ tandem periplasmic-domain protein. An initial 5-μL injection was followed by twenty 10-μL injections at 15°C, with four minutes between each injection and spinning at 340 RPM. Binding signatures were analyzed using the MicroCal Analysis software and the data were fit using a single-site model.

## Acknowledgements

This work was supported in part by grant 1R01AI148679 from the National Institute of Allergy and Infectious Disease, National Institutes of Health. The authors gratefully acknowledge Shae Padrick for advice on fluorescence anisotropy experiments and Srinivas Somarowthu for assistance in working with ^32^P-ATP.

## Notes

### Competing Interest Statement

The authors have declared no competing interest.

### Summary of Updates

Addition of new experiments showing untagged VanS protein functions similarly to the His-tagged protein; alter focus to concentrate solely on type-B resistance proteins.

