## Supplemental figures and data for "The VanS sensor histidine kinase from type-B VRE recognizes vancomycin directly"

### This PDF file includes:

***Fig. S1.*** *VanS$_B$ nanodisc preparations are competent for phosphotransfer to VanR* ... S2
***Fig. S2.*** *Vancomycin stimulation of VanS$_B$ activity: $^{32}$P-ATP assay* ... S3
***Fig. S3.*** *A VanS$_B$ construct lacking a His-tag behaves similarly to tagged protein* ... S4
***Fig. S4.*** *VanS$_B$ periplasmic domain modeling and purification* ... S5
***Fig. S5.*** *Novel AF488-based fluorescence anisotropy probe* ... S6
***Fig. S6.*** *ITC control experiments* ... S7
***Fig. S7.*** *Pairwise sequence alignment of VanS$_A$ and VanS$_B$* ... S8
***Fig. S8.*** *Lengths of the sensor domains in different VanS proteins* ... S9
***Fig. S9.*** *VanR$_B$ is partially phosphorylated in E. coli* ... S10

***Detailed Methods.***
*Autophosphorylation assay using ATPγS* ... S11
*Dephosphorylation assay* ... S11
*Liquid-chromatography mass spectrometry (LC-MS)* ... S12
*Photolabeling reactions* ... S12
*Modeling of the VanS$_B$ periplasmic domain* ... S12
*Preparation of AlexaFluor488-labeled vancomycin* ... S13
***Table S1.*** *Primers used to prepare expression constructs.* ... S14
***Supporting Information References*** ... S15

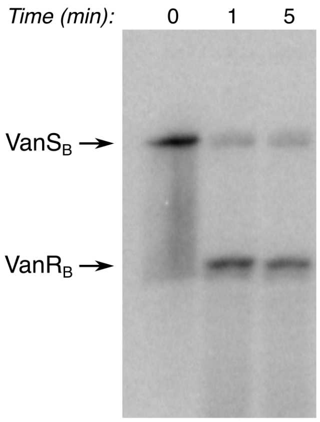


***Figure S1.*** *VanS$_B$ nanodisc preparations are competent for phosphotransfer to VanR$_B$.* Purified VanS$_B$ protein was incorporated into nanodiscs and allowed to autophosphorylate in the presence of $^{32}$P-labeled ATP. Nucleotide was removed and purified VanR$_B$ was added; aliquots were removed at the times indicated, the reaction was stopped by addition of EDTA, and samples were analyzed by autoradiography.

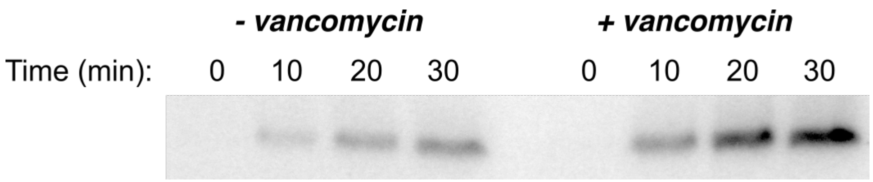


***Figure S2****. Stimulation of VanS$_B$ autophosphorylation activity by vancomycin—$^{32}$P-ATP assay.* VanS$_B$ in nanodiscs was incubated with a mixture of $^{32}$P-labelled ATP and cold ATP, ± 100 μM vancomycin, for 0, 10, 20, and 30 minutes. The reactions were subjected to SDS-PAGE and dried onto a nitrocellulose membrane, and the membrane was exposed overnight. The stimulatory effect of vancomycin on the autophosphorylation activity of VanS$_B$ is still observed when the native substrate is used.

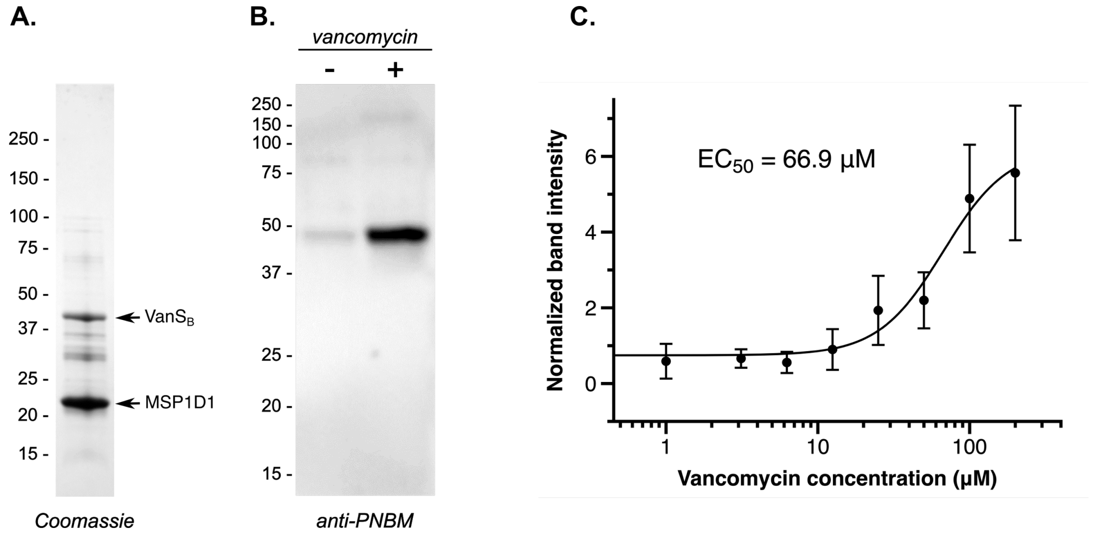


***Figure S3.*** *Full-length VanS$_B$ lacking a C-terminal His$_6$ tag behaves comparably to the tagged enzyme.* (*A*) Coomassie-stained gel showing the nanodisc preparation for the tagless VanS$_B$ protein. (*B*) Representative Western blot showing the autophosphorylation activity of the tagless VanS$_B$ nanodisc preparation ± 100 µM vancomycin. (*C*) Dose-response curve showing the dependence of VanS$_B$ autophosphorylation upon vancomycin; error bars correspond to standard deviations reflecting two independent triplicate measurements. Vancomycin stimulates the activity of the tagless VanS$_B$ protein to a similar degree as that seen with the C-terminally His-tagged protein.

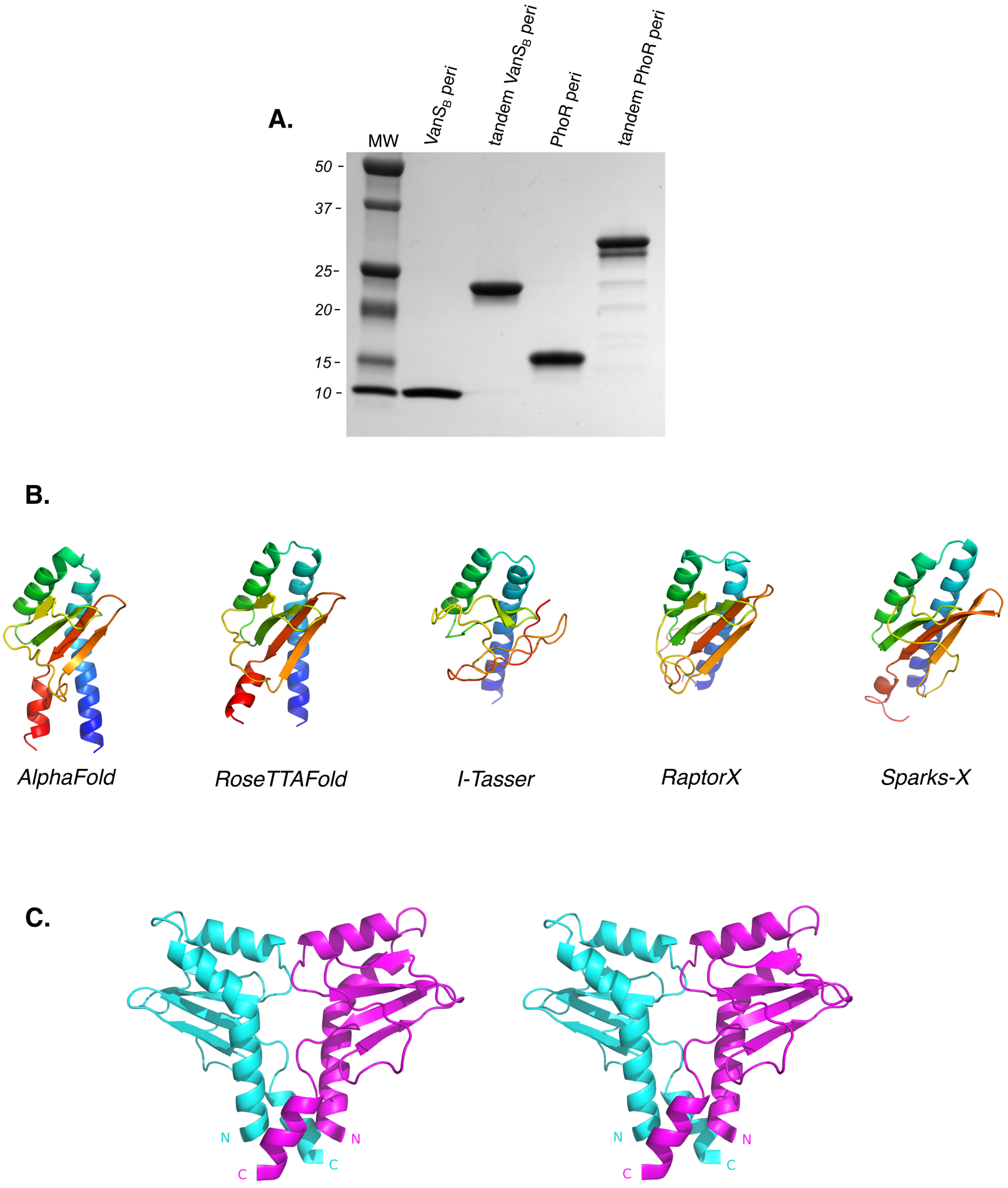


***Figure S4.*** *The VanSB periplasmic sensor domain.* A) Purified periplasmic sensor-domain constructs. Shown is a Coomassie-stained SDS PAGE gel; each lane contains 5 µg of the purified periplasmic-domain construct indicated. Molecular-weight markers are shown at left. B) Models of the $VanS_B$ periiplasmic domain produced by selected programs. Each model is colored using a rainbow scheme, in which the color gradually changes from blue at the N-terminus to red at the C-terminus. C) Divergent stereo view of the AlphaFold model of the $VanS_B$ periplasmic domain dimer. Shown are residues 35-135, with the two protomers being colored magenta and cyan. Positions of the N- and C-termini are indicated.

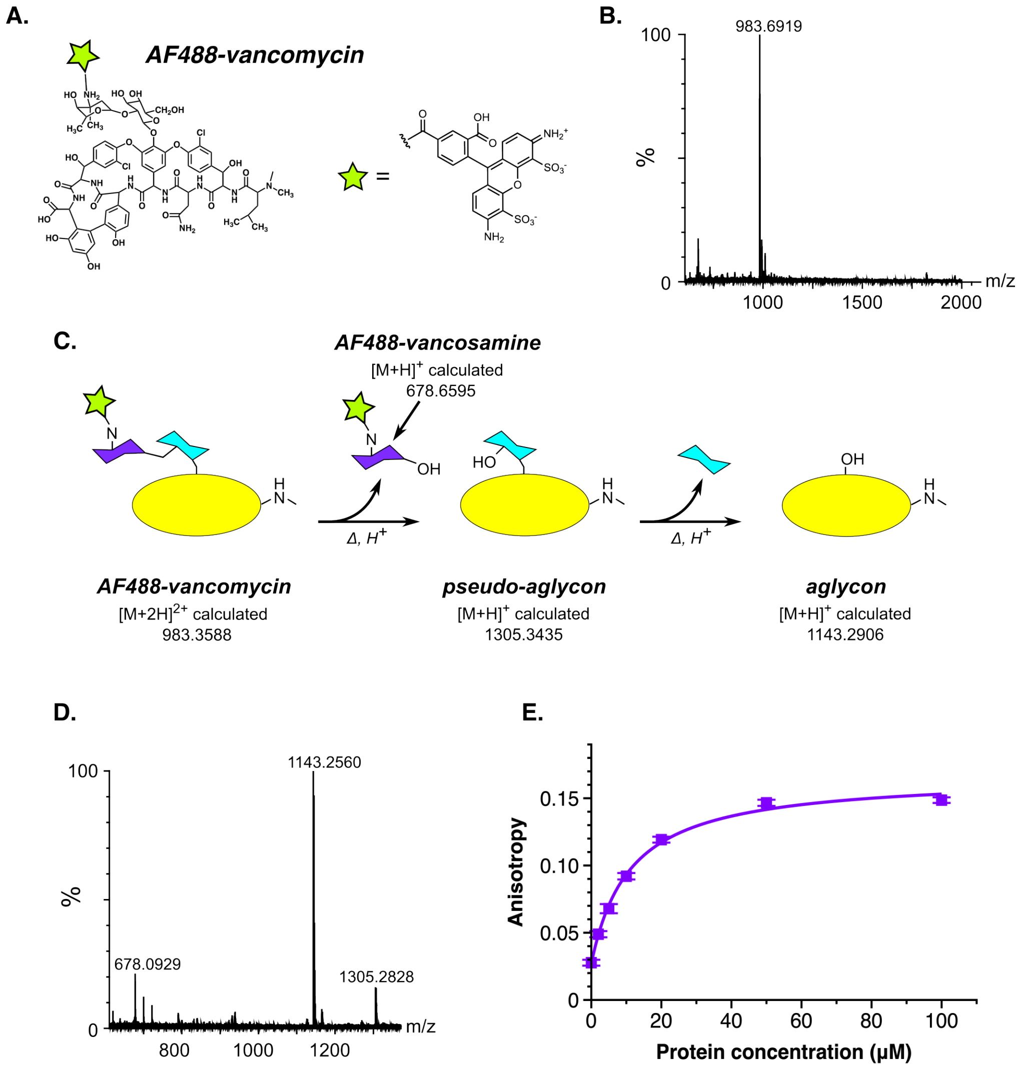


***Figure S5.*** *A new AF488-based fluorescence-anisotropy probe.* A) Structure of the new AF488-vancomycin fluorescence-anisotropy probe. B) Confirmation of mass of the probe (calculated mass, 983.3588; observed, 983.6919). C) Scheme for determining the site of attachment of the AF488 fluorophore. There are two amines in vancomycin that can react with the NHS-AF488: A primary amine in the vancosamine sugar and a secondary amine at the N-terminus of the peptide. Acid hydrolysis will release the sugars, making it possible to use mass spectrometry to determine whether the dye is attached to the sugar or the aglycon. D) Mass spectrum of the acid hydrolysis products, showing that the dye is attached to the vancosamine sugar. E) Binding of the tandem $VanS_B$ sensor domain to AF488-containing probe, as measuring by change in fluorescence anisotropy. Fit to a single binding-site model is shown; estimated $K_D$ value is 11.0 μM.

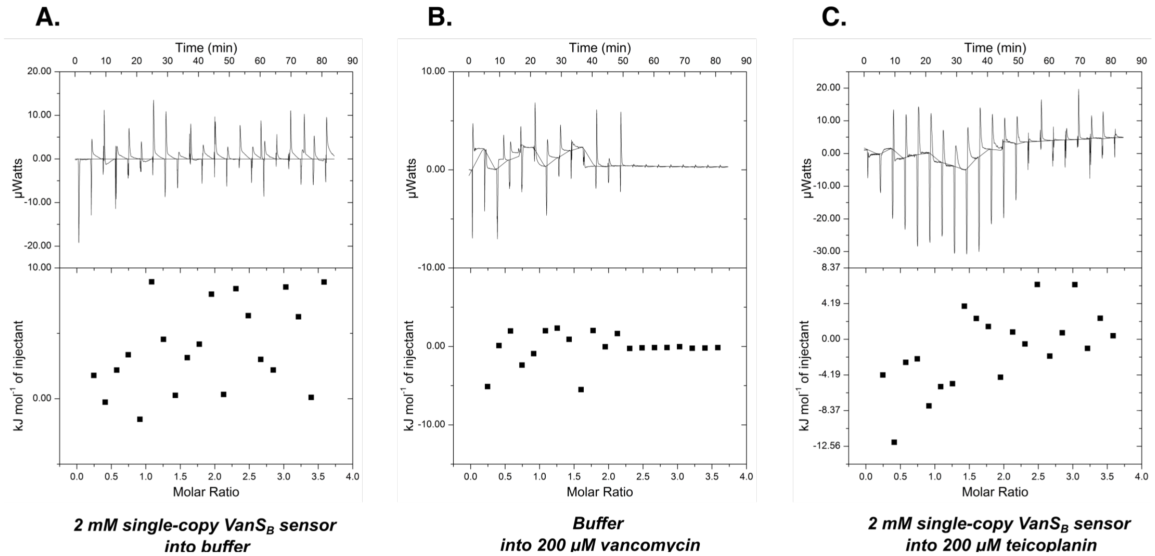


***Figure S6.*** *ITC control experiments.* (A) Injection of 2 mM $VanS_B$ sensor domain into 20 mM Bicine, pH 7.5. (B) Injection of 20 mM Bicine, pH 7.5 into 200 μM vancomycin. (C) Injection of 2 mM $VanS_B$ sensor domain into 200 μM teicoplanin.

```
VanSA    1 MVIKLKNKKNDYSKLERK-------LYMYIVAIVVVAIV-------FVLYIRSMIRG---KLGDWILSILENKYDLNHLDAMKLYQYSIR----------
                         :|||        |  || :::| :        || | |:|      |    :: :::|   |:  :   |: |
VanSB    1 --------------MERKGIFIKVFSYTIIVLLLLVGVTATLFAQQFVSYFRAMEAQQTVKSYQPLVELIQNSDRLDMQEVAGLFHYNNQSFEFYIEDKE
                                 <------ TM1 ------>

VanSA   71 ----------------------------------------------NNIDIFIYVAIVISILILCRVMLSKFAKYFDEINTGIDVLIQNEDKQ-------
                                                           |   : :||::   :||  : ::      :: | |  |  : :|
VanSB   87 GSVLYATPNADTSNSVRPDFLYVVHRDDNISIVAQSKAGVGLLYQGLTIRGIVMIAIMVVFSLLCAYIFAR------QMTTPIKALADSANKMANLKEVP
                                                           <------ TM2 ------>

                                                                ★
VanSA  121 --IELSAEMDVMEQKLNT----LKRTLEKREQDAKLAEQRKNDVVMYL----AHDIKTPLTSIIGYL-SLLDEAPDMPVDQKAKYVHITLDKAYRLEQLI
             :|   |:  :   :::    || |: : | |    |    :   |     :|::|||: ::   |  :|:   |      :||:   :   |  : |
VanSB  181 PPLERKDELGALAHDMHSMYIRLKETIARLE-DEIAREHELEETQRYFFAASHELKTPIAAVSVLLEGMLENIGD--YKDHSKYLRECIKMMDRQGKTI
                                                                ★

VanSA  210 DEFFEITRYNLQTITLTKTHIDLYYMLVQMTDEFYPQLSAHGKQAVIHAPEDLTVSGDPDKLARVFNNILKNAAAYSEDNSIIDITAGLSGDVVSIEFKN
            |  |:   |   |      :|:   : ::  :|     |: :: |   |    |  ||  : :  :|:: ||   :     : | :    :   :   |
VanSB  278 SEILELVSLNDGRIVPIAEPLDIGRTVAELLPDFQTLAEANNQRFVTDIPAGQIVLSDPKLIQKALSNVILNAVQNTPQGGEVRIWSEPGAEKYRLSVLN

VanSA  310 TG-SIPKDKLAAIFEKFYRLDNARSSDTGGAGLGLAIAKEIIVQHGGQIYAESNDNYTTFRVELPAMPDLVDKRRS    384
            |  |    |: :|  |||:| |||  :| :|||||| :: :     |   |:  :   |  ::||    |
VanSB  378 MGVHIDDTALSKLFIPFYRIDQARSRKSGRSGLGLAIVQKTLDAMSLQYALENTSDGVLFWLDLPPTSTL------    447
```

***Figure S7.*** *Pairwise sequence alignment of* $VanS_A$ *and* $VanS_B$*.* Alignment was calculated using EMBOSS-Needle (1). Vertical lines mark identities, while double dots show residues of similar character. The predicted positions of the two transmembrane helices of $VanS_B$ are highlighted in red; the periplasmic sensor domain lies between these two helices. The histidine that is the target of autophosphorylation is indicated with a star.

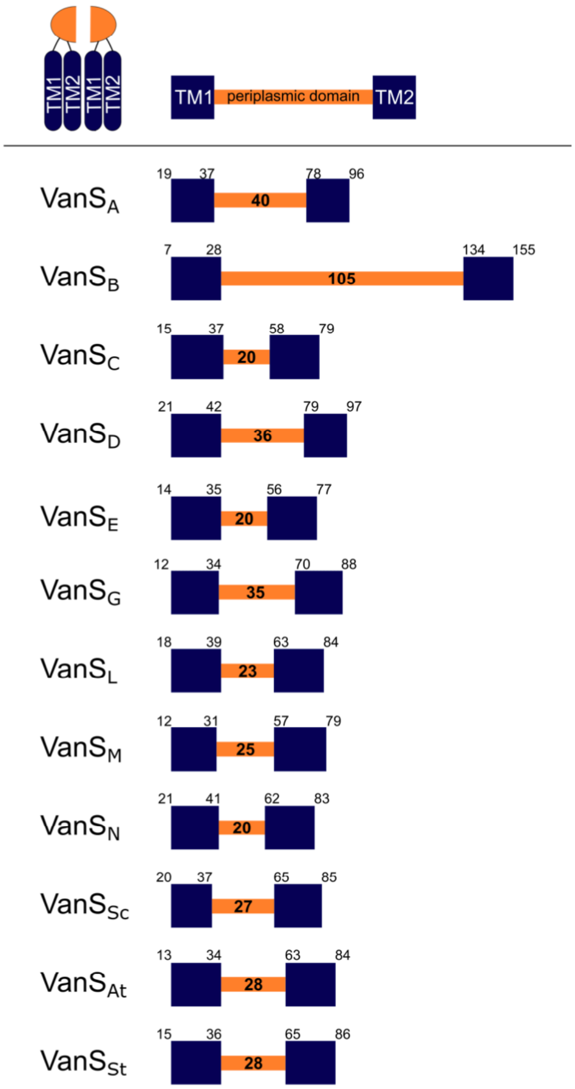


***Figure S8****. Lengths of the sensor domains in different VanS proteins.* (*Top*) Cartoon representing the generic structure of the sensing domain of a VanS protein, in which the periplasmic region is located between transmembrane helix 1 (TM1) and transmembrane helix 2 (TM2). Two such domains are expected to associated in a functional dimer. (*Bottom*) Schematic representations for the sensor domains of individual VanS proteins. The positions of the transmembrane helices were determined using CCTOP (2), and are noted in each cartoon. The lengths of different VanS periplasmic sensor domains vary in length from 20-105 amino acids; the number of residues in each is indicated in bold. Accession numbers for the sequences used: $VanS_A$, Q06240.1; $VanS_B$, Q47745.1; $VanS_C$, AAF86642.1; $VanS_D$, AAD42181.1; $VanS_E$, AAL27446.1; $VanS_G$, AAQ16269.1; $VanS_L$, ABX54692.1; $VanS_M$, ACL82958.1; $VanS_N$, AEP40504.1. Sc, At, and St refer to the VanS proteins from *Steptomyces coelicolor*, *Actinoplanes teichomyceticus*, and *Streptomyces toyocaensis*, accession numbers TYP17068, TWG11274, and AAM80542, respectively.

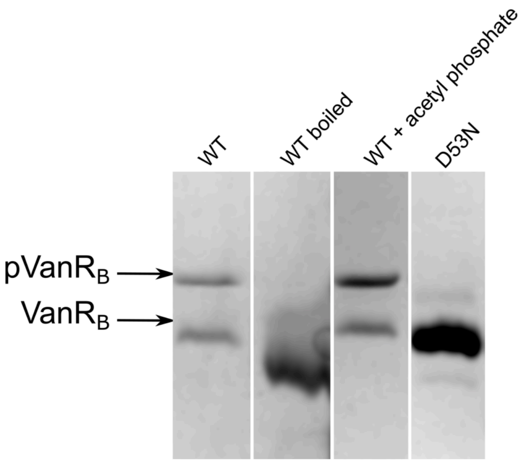


***Figure S9.*** *VanR$_B$ is partially phosphorylated in E. coli.* Coomassie-stained Phos-tag™ gel showing that VanR$_B$ as isolated from *E. coli* is partially phosphorylated. From left to right: 1) Wild-type (WT) VanR$_B$ runs as two distinct bands; 2) Boiling WT VanR$_B$ converts the two bands to a single species. Boiling is expected to remove the labile phosphoryl group; 3) WT VanR$_B$ was phosphorylated with the small-molecule phosphoryl donor acetyl phosphate, which increases the intensity of the upper band; 4) The nonphosphorylatable VanR$_B$ D53N mutant exhibits no upper band, consistent with that band corresponding to phospho-VanR$_B$.

***Detailed Methods***

***Autophosphorylation assay using ATPγS.*** Autophosphorylation assays used the following stock solutions: 5X reaction buffer (250 mM KCl, 50 mM $MgCl_2$, 250 mM Tris pH=7.4), 10 mM ATPγS (Abcam ab138911) in ultrapure water, 1 mM vancomycin in ND buffer (20 mM Tris pH 7.4, 150 mM NaCl), 0.5 M EDTA pH 8, and 50 mM p-nitrobenzyl mesylate (PNBM; Abcam ab138910) in 100% dimethyl sulfoxide.

To determine the effect of vancomycin on autophosphorylation activity, two separate master-mix reactions were created, of 66 µL each. Each reaction contained 1x reaction buffer, ATPγS at a final concentration of 1 mM, and VanS-containing nanodiscs at a final concentration of 0.3-0.5 µM. One reaction contained vancomycin at a final concentration of 100 µM and the other contained an equivalent volume of ND buffer to ensure the salt concentration was consistent. At each time point, 15 µL was transferred from the reaction to a designated tube that contained 3 µL of 0.5 M EDTA pH 8 to quench the reaction. After all of the time points had been taken, 2 µL of PNBM was added to each tube and the mixture was incubated at room temperature for 1 hour. The samples were loaded onto a Bio-Rad 12% precast gel and run at 200 V with cold running buffer at 4°C, and subsequently transferred onto PVDF membrane for 1 hour at 100 V. The membrane was rocked in blocking buffer (5% milk in Tris-buffered saline, 0.1% Tween 20 (TBST)) for 1 hour, rocked in anti-PNBM (Abcam ab92570) at 1:5000 in 1% milk-TBST for 1 hour, washed in TBST 6 × 8min, rocked in anti-GAR (Jackson ImmunoResearch HRP-GAR IgG 111-035-003) at 1:1000 in 1% milk-TBST for 1 hour, and washed in TBST 6 × 8min. The membranes were rocked in enhanced chemiluminescence detection reagent for 1 min prior to exposing the membranes using a Syngene imager. Blots were stripped and probed with HRP-conjugated anti-His antibody at 1:20,000 dilution in 1% milk TBST. The band intensities and background intensities were quantified by ImageJ and the band intensities for each time point were background-corrected. To determine the fold-stimulation by vancomycin, bands representing autophosphorylation in the presence of vancomycin were normalized to corresponding time points representing autophosphorylation in the absence of vancomycin.

A slight variation to the above protocol was introduced in order to determine the $EC_{50}$ of vancomycin. Multiple individual 15-µL autophosphorylation reactions were initiated in the presence of varying concentrations of vancomycin. The start of the reactions was staggered by 30 seconds, and each reaction was allowed to proceed for 30 minutes, at which point 3 µL of EDTA was added to quench.

***Dephosphorylation assays.*** Dephosphorylation reactions consisted of 1x reaction buffer, $VanS_B$-containing nanodiscs at 0.5 µM, and $VanR_B$ at 1 µM. $VanR_B$ purified from *E. coli* was found to be partially phosphorylated and was used as-is, without additional phosphorylation (Figure S8). Dephosphorylation reactions were quenched by addition of SDS-PAGE loading buffer and samples were loaded onto Phos-tag gels (Fujifilm Cat# 195-17991). Gels were run in cold, fresh running buffer at 200 V for approximately 2-2.5 hours at 4°C, and then stained with Coomassie Brilliant Blue.

***Liquid-chromatography mass spectrometry.*** Molecules were analyzed on a Waters Acquity I-Class UPLC system coupled to a Synapt G2Si HDMS mass spectrometer in positive ion mode with a heated electrospray ionization (ESI) source in a Z-spray configuration. For proteins, LC separation was performed on a Waters Acquity UPLC Protein BEH $C_4$ 1.7 µm 2.1 x 50mm column maintained at 40 °C; a 0.2 mL/min gradient of 80/20 to 30/70 A/B in 20 min was used, followed by washing and reconditioning the column. For vancomycin, LC separation was performed on a Waters Acquity UPLC BEH $C_{18}$ 1.7 µm 2.1 x 50 mm column using an 0.6 ml/min gradient of 95/5 to 15/85 A/B over the course of four minutes. Eluent A is 0.1% v/v formic acid in water and B is 0.1% v/v formic acid in acetonitrile. Conditions on the mass spectrometer were as follows: capillary voltage 0.5 kV, sampling cone 40 V, source offset 80 V, source 120 °C, desolvation 250 °C, cone gas 0 L/h, desolvation gas 1000 L/h and nebulizer 6.5 bar. The analyzer was operated in resolution mode and low energy data was collected between 100 and 2000 Da at 0.2 sec scan time. For vanocmycin, $MS^e$ data was collected using a 20-40V ramp trap collision energy. Masses were extracted from the TOF MS TICs using a 0.005 Da abs width. Protein ESI data was deconvoluted using MaxEnt1 in Masslynx 4.1 (Waters Corporation).

***Photolabeling reactions.*** 50-µL reactions containing 20 µM protein and 50 µM photoprobe were prepared in 20 mM Tris pH 7.5, 100 mM NaCl. The reactions were placed into glass depression spot plates, which were placed atop a heavy aluminum plate packed in ice, to keep reactions cold and to limit evaporation. Samples were irradiated at 365 nm using a Stratagene Stratalinker 2400 UV Crosslinker for the times specified. For competition experiments with vancomycin, samples were irradiated for 2 minutes. All samples were diluted 1:50 for subsequent analysis by mass spectrometry and Western blotting.

For Western-blot analysis, 100 ng of protein were loaded onto a 12% SDS-PAGE gel and electrophoresed at 180 V. While the gel was running, a 0.2 µm PVDF membrane (Cytiva #10600021) was prepared by soaking in 100% methanol for one minute then in water for two minutes. The membrane was then equilibrated in transfer buffer for 15 minutes (25 mM Tris pH 8.3, 192 mM glycine, 15% methanol). Proteins were transferred from the gel to the membrane for one hour at 100 V, after which the membrane was rocked in blocking buffer (5% milk in 20 mM Tris pH 7.6, 150 mM NaCl, 0.1% Tween-20 (TBST)) for 30 minutes. The membrane was then rocked overnight at 4°C in a solution containing a sheep anti-vancomycin polyclonal antibody (Bio-Rad # 9520-0004) diluted 1:1000 in blocking buffer. The following day, the membrane was washed in TBST 3 x 10min and then rocked at room temperature for one hour in solution containing a rabbit anti-sheep HRP-conjugated secondary antibody (Invitrogen # 31480), diluted 1:5000 in blocking buffer. The membrane was then washed in TBST 3 x 10 min and placed in peroxidase substrate solution (Pierce #32209) for 1 minute before chemiluminescent detection.

***Modeling of the VanS$_B$ periplasmic sensor domain.*** The sequence of the VanS$_B$ periplasmic domain (residues 31-132, Uniprot Q47745) was used for protein threading (template-based) experiments, using I-TASSER, RaptorX, Phyre2, and SparksX (3-6). All the threading approaches used gave predicted structures with similar folds; selected examples are shown in Figure S4. One of the templates chosen most frequently by the different threading methods was the crystal structure of the periplasmic sensor domain of PhoR from *B. subtilis* (PDB ID 3CWF), which prompted our use of this protein as a negative control that has a similar fold but a different function. For *ab initio* (template-independent) modeling studies, we used Quark, RoseTTAFold, and AlphaFold2 (7-9). In particular, AlphaFold2 and RoseTTAFold produced very similar structures (Fig. S4), with an RMSD for all C$\alpha$ positions of 1.63 Å. Given that sensor histidine kinases are widely understood to be obligate dimers, we used AlphaFold2 to generate a model for the dimer, which is shown in panel (C) of Figure S4.

***Preparation of AF488-labeled vancomycin.*** Vancomycin was dissolved at a concentration of 25 mg/mL in 90 mM sodium phosphate, pH 8. One mg of AF-488 NHS ester (BroadPharm, cat. no. BP-24307) was dissolved in 50 µL of DMSO. 150 µL of the vancomycin solution (3.75 mg, 2.6 µmol) was added to the AF-488 NHS ester (1 mg, 1.6 µmol) and incubated at room temperature, protected from light, for 2 hour. The reaction mixture was then frozen. After thawing, it was diluted 1:10 with water, after which DMSO was added to a final concentration of 20% (v/v). The material was then purified by reverse-phase chromatography on a 1 x 20 cm Ultrasphere 5ODS column (Hichrom, Leicestershire, UK), using gradient from 5% A to 100% B (solvent A = 0.25% formic acid in water, solvent B = 0.25% formic acid in acetonitrile). Two peaks having identical masses were isolated, corresponding to the addition of the fluorophore to either the vancosamine sugar or the antibiotic N-terminus ($[M+2H]^{2+}$ calculated, 983.3588; observed, 983.6919). To distinguish these two species, the sugars were removed from the aglycon macrocycle by acid hydrolysis, after which the mass of the aglycon was determined by mass spectrometry (10, 11; see Figure S5). We were able to cleanly isolate the species having the fluorophore attached to the vancosamine sugar (~95% pure); however, we were unable to isolate clean fractions containing the species labeled at the antibiotic's N-terminus, and therefore did not use this species for binding experiments.

| Primer | Primer name | Primer Sequence |
|---|---|---|
| **1** | VanR$_B$_pETCH_F | 5′-TTTTTTTTCCATGGCGATACGAATTCTACTTGTCGA-3′ |
| **2** | VanR$_B$_pETCH_R | 5′-TTTTTTTTTTCCCGGGTAATGATTCCTCCAATCGGTAAC-3′ |
| **3** | VanR$_B$_D53N_F | 5′-GTTATTCTTAATATTATGCTGCCCGGTATGAATGGGCATGAA-3′ |
| **4** | VanR$_B$_D53N_R | 5′-CAGCATAATATTAAGAATAACCAGTTGATAGGTGTTTTCATAGAACTTG-3′ |
| **5** | VanS$_B$_pETCH_F | 5′-TTTTTTTTTTCCATGGAACGCAAAGGCATCTTCATC-3′ |
| **6** | VanS$_B$_pETCH_R | 5′-TTTTTTTTTCCCGGGCAGCGTTGAGGTCGGCG-3′ |
| **7** | VanS$_B$_pETHSUL_F | 5′-CCGCGAACAGATTGGTGGCGGTATGGAACGCAAAGGCATCTTCAT-3′ |
| **8** | VanS$_B$_pETHSUL_R | 5′-CTTCTCGAGGAGAGTTTAGACGATTACAGCGTTGAGGTCGGCGGC-3′ |
| **9** | cVanS$_B$_pETCH_F | 5′-TTTTTTTTTTCCATGGCCACGCCGATCAAAGCCCT-3′ |
| **10** | cVanS$_B$_pETCH_R | 5′-TTTTTTTTTTCCCGGGCAGCGTTGAGGTCGGCGG-3′ |
| **11** | VanS$_B$_periplasmic_STOP_F | 5′-TCAAGGTTAATGCGGTGGTAGTCAGCAGTTCGTTTCATATTTCCGTGCC-3′ |
| **12** | VanS$_B$_periplasmic_STOP_R | 5′-CCACCGCATTAACCTTGATACAGCAGGCCAACACCGGCTTTGC-3′ |
| **13** | PhoR_Bs_fwd | 5′-GAGAACCTGTACTTCCAGATGGAAACATCTGATCAAAGGAAAGCAG-3′ |
| **14** | PhoR_Bs_rev | 5′-GAAGCTTATTTAATTACCTGCATTACAATTCGCCTTTTAAGCCGTCGC-3′ |
| **15** | PhoR_Bs_int_fwd | 5′-GAATTGGGCTCTGGCGGTTCTGAAACATCTGATCAAAGGAAAGCAG-3′ |
| **16** | PhoR_Bs_int_rev | 5′-GTTTCAGAACCGCCAGAGCCCAATTCGCCTTTTAAGCCGTCGC-3′ |

***Table S1**. Primers used to prepare expression constructs.*

**Supporting Information References**
